# Hepatic lipid-associated macrophages mediate the beneficial effects of bariatric surgery against MASH

**DOI:** 10.1101/2023.06.11.544503

**Authors:** Gavin Fredrickson, Kira Florczak, Fanta Barrow, Katrina Dietsche, Haiguang Wang, Preethy Parthiban, Rawan Almutlaq, Oyedele Adeyi, Adam Herman, Alessandro Bartolomucci, Christopher Staley, Cyrus Jahansouz, Jesse Williams, Douglas G. Mashek, Sayeed Ikramuddin, Xavier S. Revelo

**Affiliations:** Department of Integrative Biology & Physiology, University of Minnesota, Minneapolis, MN 55455, USA; Department of Laboratory Medicine and Pathology, University of Minnesota, Minneapolis, MN 55455, USA; Department of Surgery, University of Minnesota, Minneapolis, MN 55455, USA; Center for Immunology, University of Minnesota, Minneapolis, MN 55455, USA; Department of Biochemistry, Molecular Biology, and Biophysics, University of Minnesota, Minneapolis, MN 55455, USA; Division of Diabetes, Endocrinology, and Metabolism, Department of Medicine, University of Minnesota, Minneapolis, MN, USA

## Abstract

For patients with obesity and metabolic syndrome, bariatric procedures such as vertical sleeve gastrectomy (VSG) have a clear benefit in ameliorating metabolic dysfunction-associated steatohepatitis (MASH). While the effects of bariatric surgeries have been mainly attributed to nutrient restriction and malabsorption, whether immuno-modulatory mechanisms are involved remains unclear. Here we report that VSG ameliorates MASH progression in a weight loss-independent manner. Single-cell RNA sequencing revealed that hepatic lipid-associated macrophages (LAMs) expressing the triggering receptor expressed on myeloid cells 2 (TREM2) increase their lysosomal activity and repress inflammation in response to VSG. Remarkably, TREM2 deficiency in mice ablates the reparative effects of VSG, suggesting that TREM2 is required for MASH resolution. Mechanistically, TREM2 prevents the inflammatory activation of macrophages and is required for their efferocytotic function. Overall, our findings indicate that bariatric surgery improves MASH through a reparative process driven by hepatic LAMs, providing insights into the mechanisms of disease reversal that may result in new therapies and improved surgical interventions.

## MAIN

Metabolic dysfunction-associated steatotic liver disease (MASLD) is estimated to affect 30% of the population and is one of the leading causes of abnormal liver function^1^. MASLD is highly associated with obesity and covers a wide spectrum of liver pathology ranging from simple lipid accumulation to the more serious metabolic dysfunction-associated steatohepatitis (MASH), characterized by inflammation, hepatocellular injury, and fibrosis^2, 3^. Hepatic inflammation is primarily driven by liver macrophages and is a critical component in the initiation and progression of MASH^3, 4^. Although lifestyle and dietary intervention can result in moderate weight loss with concomitant improvements in comorbidities, these effects are transient, and weight regain is common^5^. Bariatric procedures such as vertical sleeve gastrectomy (VSG) are the most successful and effective treatment options for obese adults^6^ and are effective at ameliorating the progression of MASH^7, 8, 9^. While changes in body weight have a profound metabolic impact, bariatric surgery often results in improved insulin sensitivity before any substantial weight loss, indicative of weight-loss-independent effects^10, 11^. Several mechanisms of weight-loss-independent improvements in metabolism due to bariatric surgery have been proposed including bile acid signaling^12^, gut hormones^13^, intestinal tissue reprogramming^14^, changes in the gut microbiome^15, 16^, and HIF2α-mediated iron absorption^17^. However, whether bariatric surgery ameliorates the progression of MASH through immunomodulatory mechanisms has not been investigated.

Recent research has shown that MASH is associated with the emergence of macrophages with a lipid-associated signature, termed hepatic lipid-associated macrophages (LAMs), that express the triggering receptor expressed on myeloid cells-2 (TREM2)^18, 19^. Hepatic LAMs are conserved in mice and humans, correlate with disease severity, preferentially localize to steatotic regions, and are highly responsive to dietary intervention^19, 20^. In MASH, TREM2 expression in LAMs is required for effective efferocytosis of apoptotic hepatocytes^21^. However, prolonged hypernutrition and hepatic inflammation promote the shedding of TREM2 from the cell surface of LAMs leading to an ineffective clearance of dying hepatocytes^21^. Cleavage of TREM2 increases soluble TREM2 (sTREM2) in the circulation, making it a promising biomarker of MASH severity^22^. TREM2 is also required for adequate metabolic coordination between macrophages and hepatocytes, lipid handling, and extracellular matrix remodeling^21, 22, 23^. In a mouse model of hepatotoxic injury, TREM2 deficiency exacerbated inflammation-associated injury through enhanced toll-like receptor signaling, in agreement with its anti-inflammatory function^24^. Given the key roles of TREM2 in the diseased liver, mounting evidence has shown that TREM2 deficiency worsens most aspects of MASLD and MASH^21, 22, 23, 24, 25^. As TREM2 is a central signaling hub regulating macrophage function, therapeutic efforts have been made to stimulate the TREM2 active domain or block its shedding in neurodegenerative disease^26^. Despite the evidence suggesting that LAMs are protective, the mechanisms promoting their restorative function in MASH are poorly understood.

In the current study, we found that VSG ameliorates the progression of MASH and results in profound effects on the macrophage transcriptomic profile in a weight loss-independent manner. Remarkably, the newly described hepatic lipid-associated macrophages (LAMs) upregulate their reparative, lipid-quenching, and anti-inflammatory programs in response to bariatric surgery. Most notably, TREM2 deficiency prevented VSG-induced MASH reversal, suggesting that TREM2 is essential for the beneficial effects of bariatric surgery in the liver. Mechanistically, TREM2 prevents the production of inflammatory cytokines by macrophages and is required for adequate efferocytosis of apoptotic hepatocytes. Overall, our results indicate that bariatric surgery improves MASH through a reparative process driven by hepatic LAMs.

## RESULTS

### Vertical sleeve gastrectomy ameliorates MASH progression independent of weight loss

To study the mechanisms by which bariatric surgery ameliorates MASH, we used a validated mouse model of VSG in which approximately 80% of the lateral stomach of mice is clamped by a gastric clip and excised^15^. To induce MASH, we fed mice a high-fat high-carbohydrate (HFHC) diet that recapitulates aspects of human disease such as obesity, hepatic lipid accumulation, inflammation, and fibrosis^27^. Mice were fed the HFHC diet ad libitum for 12 weeks and then assigned to either sham (Sham AL) or VSG surgery (**Fig. 1A**) and remained on the HFHC diet for 5 weeks. Most mice survived the surgery, and we confirmed surgical anatomy by oral gavage of barium and imaging^15^. To determine weight-loss-independent effects, we included a sham group that was pair-fed to the VSG group (Sham PF) to match their caloric intake during the post-surgery period^28^. In addition, a group of mice were fed a normal chow diet (**NCD**) for the duration of the studies. Compared with Sham AL, both VSG and Sham PF mice resulted in a similar decrease in average daily food intake (**Fig. 1B**). Following the surgeries, body weight rapidly decreased in Sham AL, PF, and VSG groups due to the intervention, as previously reported^29^ (**Fig. 1C**). However, while the body weight in Sham AL mice recovered, Sham PF and VSG mice maintained their weight loss throughout the study (**Fig. 1C**). Five weeks after the surgeries, both Sham PF and VSG mice showed a similar decrease in liver weight (**Fig. 1D**). Compared with Sham AL controls, only the VSG group showed a reduction in hepatic lipid accumulation (**Fig. 1E and 1F)**, ALT/AST (**Fig. 1G**), fibrosis (**Fig. 1H and Extended Data Fig. 1B**), and NAS score (**Extended Data Fig. 1A**), suggesting that the effects of VSG are partly independent of the reduced caloric intake induced by the surgery. As decreased intestinal lipid absorption has been proposed as a mechanism of VSG actions^30^, we measured fecal lipids and found that VSG mice had increased fecal lipids compared with Sham AL mice (**Fig. 1I**). We also determined the effects of VSG on MASH progression 10 weeks after surgeries and found that the VSG-induced reductions in body and liver weight, liver triglycerides, AST, and ALT were maintained at this later timepoint (**Extended Data Fig. 1C**-G). Overall, these findings indicate that VSG results in substantial improvements in MASH in a weight-loss-independent manner.

**Figure 1.**
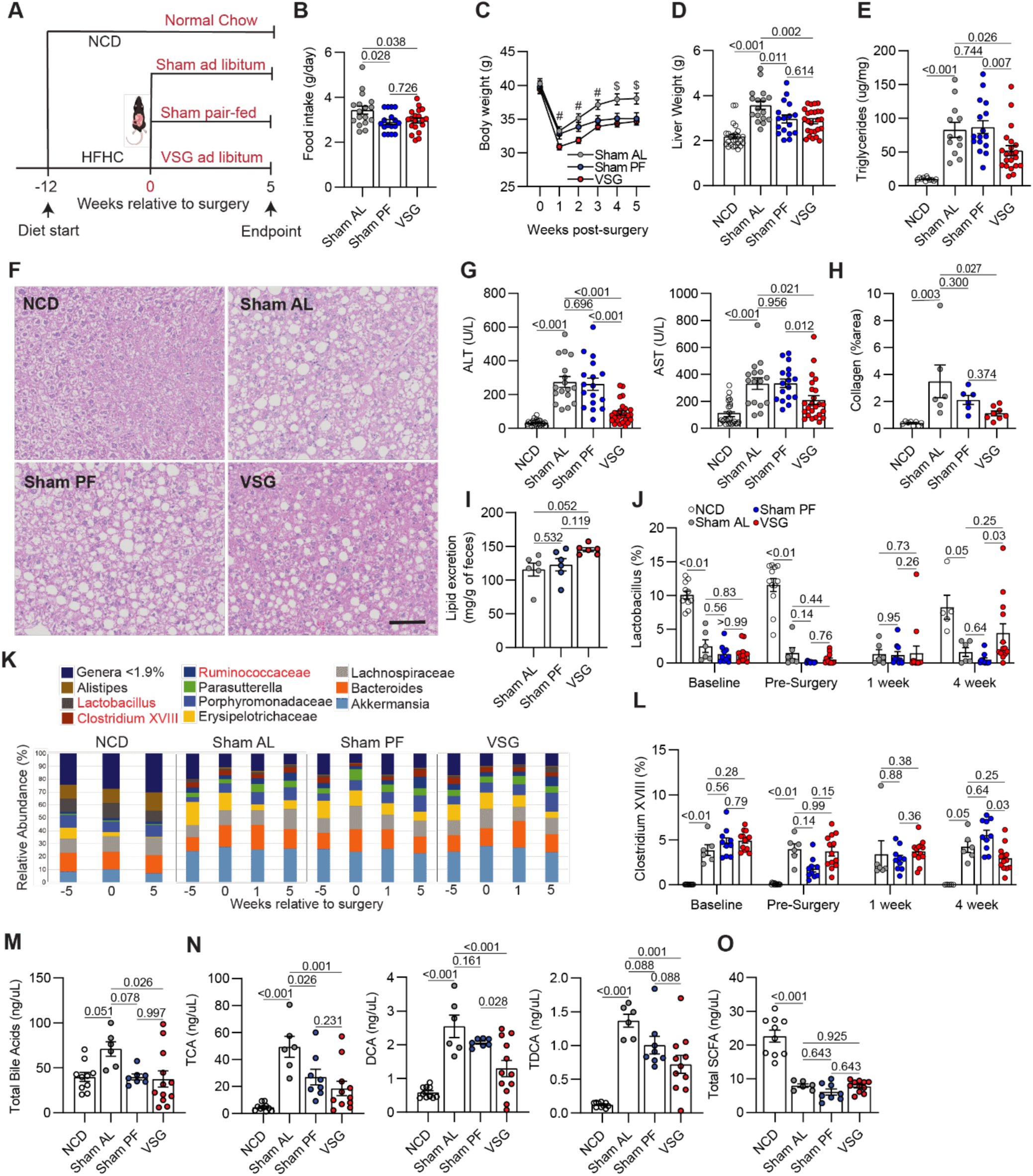
Vertical sleeve gastrectomy (VSG) ameliorates MASH progression independent of weight loss. (**A**) Experimental design, (**B**) Mean daily food intake post-surgery, (**C**) Weekly body weights post-surgery, (**D**) Liver weight, (**E**) Hepatic triglyceride content, (**F**) Representative H&E-stained liver sections, (**G**) Serum ALT (Left) and AST (Right), (**H**) Collagen area in Picro Sirius Red-stained liver sections, (**I**) Fecal lipid content, (**J**) Relative abundance of Lactobacillus, (**K**) Relative abundance of microbiota species, (**L**) Relative abundance of Clostridium XVIII, (**M**) Total portal vein bile acids, (**N**) Concentration of portal vein taurocholic acid (TCA), deoxycholic acid (DCA), and taurodeoxycholic acid (TDCA), and (**O**) Total portal vein short-chain fatty acids (SCFA) in C57BL6/J (WT) mice fed an HFHC diet for 12 weeks before assignment to Sham ad libitum (Sham AL; n = 6-17), Sham pair-fed (Sham PF; n = 6-17), or VSG (n = 6-25) surgeries. Mice were maintained on the HFHC diet for 5 weeks post-surgery. A cohort of mice without any intervention were fed a normal chow diet (NCD; n = 5-25) throughout the study. Data are biological experimental units presented as mean ± standard error of the mean (SEM). Except for the microbiota data, which were analyzed by a non-parametric Kruskal-Wallis test, data were analyzed by one-way ANOVA with Holm-Šídák multiple comparison test.

To determine if the VSG-induced improvements in MASH were associated with alterations in the gut microbiota and its metabolites, we analyzed the fecal microbial composition of NCD, Sham AL, Sham PF, and VSG mice 5 weeks after surgeries. Compared with NCD mice, HFHC feeding induced dramatic remodeling in gut microbial species, including increased abundances of *Akkermansia*, *Erysipelotrichaceae*, *Parasutterella*, *Ruminococceae*, *Clostridium XVIII*, and decreased abundances of *Lactobacillus*, and *Alistipes* (**Fig. 1K and Supplemental Table 1**). While HFHC feeding caused a substantial loss of *Lactobacillus*, regardless of surgical treatment, VSG resulted in a restoration of this genus towards levels found in NCD mice (**Fig. 1J**). In contrast, the HFHC diet increased the abundance of *Clostridium XVIII* whereas it was partially lowered following VSG (**Fig. 1L**). No differences in fecal microbiota composition between Sham AL and Sham PF mice were detected 5 weeks after the interventions (**Supplemental Table 1**). Intestinal products can translocate into the portal vein and reach the liver where they can regulate immune homeostasis and inhibit inflammation^31^. Thus, we measured the concentrations of bile acids (BA) and short-chain fatty acids (SCFA) in the portal vein blood. While HFHC feeding increased the total amount of BA, as observed in Sham AL mice, both Sham PF and VSG mice had decreased BA concentrations to levels similar to NCD controls (**Fig. 1M**). In particular, VSG and Sham PF mice had a similar decrease in taurine-conjugated cholic acid while VSG was more effective at reducing deoxycholic acid and taurodeoxycholic acid (**Fig. 1N**). Independent of surgical intervention, HFHC feeding decreased the levels of cholic acid, chenodeoxycholic acid, and hyodeoxycholic acid and increased several taurine-conjugated BA in all groups without any effects of Sham PF or VSG (**Extended Data Fig. 1H**). HFHC feeding also decreased the concentration of SCFAs in hepatic portal serum and neither VSG nor Sham PF had an impact on their concentration (**Fig. 1O and Supplemental Table 2**). Overall, these data suggest that the VSG-induced improvements in MASH are associated with a partial restoration of specific gut microbial species and BAs.

### scRNA-seq reveals profound effects of VSG on hepatic LAMs

To explore the impact of VSG on hepatic macrophages, we first determined the abundance of macrophages and Kupffer cells (KCs) subsets by flow cytometry^32^. We did not detect any differences in the number of monocyte-derived macrophages (MoMF), embryonic KCs (emKC), monocyte-derived KCs (moKC), and VSIG4^-^ macrophages between Sham and VSG groups at 5 or 10 weeks after surgeries (**Extended Data Fig. 2A**). To determine how VSG influences the function of hepatic macrophages, we profiled the gene expression of macrophages from Sham AL, Sham PF, and VSG mice using droplet-based single-cell RNA sequencing (scRNA-seq, **Fig. 2A**). Livers were perfused before immune cell isolation to remove circulatory cells^33^. We multiplexed samples using cell multiplexing oligos (CMOs) to track the sample of origin. Cells were loaded into the ports of a 10x-Genomics chip following a single-cell 3’ kit. The gene expression and CMO libraries were sequenced using a Novaseq S4 chip (2x150bp PE). After quality control, data were normalized and de-multiplexed. Monocytes and macrophages were identified using the cell ID function of Seurat and re-clustered for analysis. We profiled the gene expression of 58,904 single cells (avg. read depth 50,000, **Fig. 2B**). Unsupervised graph-based clustering was performed on the integrated dataset and cells were visualized using uniform manifold approximation and projection (UMAP). Independent of surgical groups, integrated UMAP analysis revealed 18 clusters of monocytes and macrophages, whose identity was determined based on the expression of established marker genes^18, 34^ (**Fig. 2C, Extended Data Fig. 2B, and Supplemental Table 3**).

**Figure 2.**
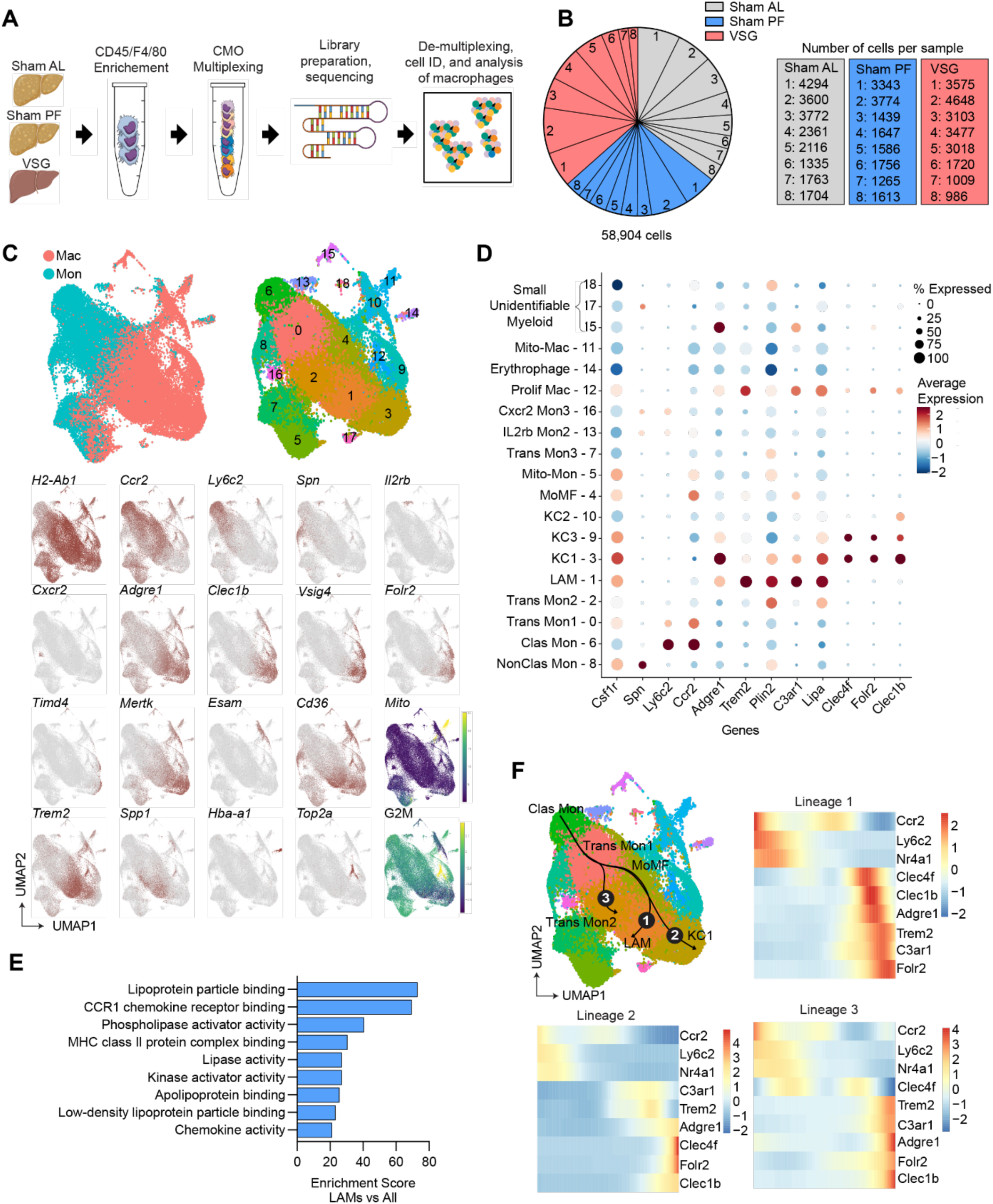
ScRNA-seq reveals profound effects of VSG on hepatic LAMs. (**A**) Experimental design, (**B**) Proportion (left) and number (right) of sequenced cells per sample, (**C**) Integrated uniform manifold approximation and projection (UMAP) analysis of all monocytes and macrophages and expression of marker genes, (**D**) Heat and dot plot of the expression and coverage of marker genes in all subsets, (**E**) Pathway analysis of upregulated differentially expressed genes (DEG) in LAMs, compared with all other subsets (Gene ontology terms, p-value < 0.1), and (**F**) Slingshot trajectory analysis (top left) and gene expression over pseudotime for trajectories 1 (top right), 2 (bottom left), and 3 (bottom right) from single-cell RNA sequencing (scRNA-seq) of hepatic macrophages and monocytes from Sham AL (n = 8), Sham PF (n = 8), and VSG (n = 8) mice 5-10 weeks post-surgeries. Differential expression testing was performed by a Wilcoxon rank-sum test.

Independent of surgical intervention, clusters 0, 2, 5, 6, 7, 8, 13, and 16 were identified as monocyte-like cells while clusters 1, 3, 4, 9, 10, 11, 12, and 14 were macrophages (**Fig. 2C**). Monocyte clusters had heterogeneous gene expression profiles indicative of their progressive stages of maturation and function. Clusters 0, 2, and 7 had dual monocyte and macrophage features, such as *Ly6c2* and *H2-Ab1*, suggesting that they were transitioning monocytes on the trajectory of becoming macrophages. Cells in cluster 6 were identified as classical monocytes based on their high expression of *Ly6c2* and *Ccr2,* which allows them to migrate in response to inflammation. Monocytes in cluster 8 were enriched in *Spn* and did not express *Ccr2* and *Ly6c2* indicating these cells were non-classical monocytes capable of patrolling. Clusters 13 and 16 corresponded to unidentifiable monocyte populations that expressed high levels of *Il2rb* and *Cxcr2,* respectively (**Fig. 2C**). We next analyzed the macrophage subsets which can broadly be divided into Kupffer cells (KCs) and MoMFs. Clusters 3, 9, and 10 were identified as KCs (KC1- 3) based on their expression of *Clec1b*, *Clec4f*, *Vsig4*, and *Folr2*. These clusters of KCs had minimal expression of *Timd4* suggesting that they are primarily moKCs^32^. Clusters 3 and 9 had a similar gene expression pattern, although cluster 3 was enriched in genes associated with efferocytosis such as *Mertk* and *Wdfy3*. Cluster 10 was enriched in *Esam* and *Cd36* and resembles a recently identified subset of pathogenic KCs termed “KC2”^35^. Among MoMFs, cells in cluster 1 were enriched for LAM genes including *Trem2*, *Spp1*, *Lipa*, and *Cd36*^36^ (**Fig. 2C, 2D and Supplemental Table 3**). Pathway analysis showed that LAMs have enriched gene programs associated with lipid metabolism such as “Lipoprotein particle binding” and “Lipase activity” (**Fig. 2E**). Cluster 4 was composed of MoMFs and transitioning monocytes that could not be further characterized based on their differential gene expression. Cells in cluster 14 were enriched in hemoglobin genes *Hba-a1* and *Hba-a2* which are highly expressed by erythrophages. Cluster 12 had a high G2M score and was enriched with *Top2a*, typical of proliferating macrophages (**Fig. 2C, 2D and Supplemental Table 3**). To gain insight into the differentiation of macrophage subsets, we performed a slingshot trajectory analysis^37^ of our integrated dataset. Although the origin (ontogeny) and cellular turnover of hepatic LAMs is unknown, recent work suggests that recruited MoMFs give rise to hepatic LAMs^22, 38^. Using monocytes as the origin, we found three primary trajectories by which newly recruited monocytes differentiate into transitioning monocytes and then either differentiate into LAMs, KC1s, or Trans Mon2s (**Fig. 2F and Extended Data Fig. 2C**). These data suggest that LAMs are derived from classical monocytes and are consistent with recent studies that have used single cell transcriptomics to reveal a previously unappreciated heterogeneity in hepatic macrophages in the MASH liver^18, 39^.

Following cluster identification and trajectory analysis, we demultiplexed the samples based on their experimental group and determined the relative abundance of each cluster. There were no differences in the proportion and number of the major myeloid clusters 5 weeks post-VSG (**Fig. 3A**). To investigate the effects of VSG on the transcriptome of hepatic macrophages, we performed differential gene expression analysis between Sham AL, Sham PF, and VSG groups and found distinct expression patterns in macrophage clusters analyzed in bulk (**Extended Data Fig. 3A**) and per cluster (**Supplemental Table 4**). Given the reparative functions of LAMs in MASH, we focused our subsequent analysis on these cells. First, we quantified the average single-cell expression of the *Trem2* gene in the LAM cluster and found no differences between groups (**Fig. 3B**). Despite no effect on the abundance of LAMs and *Trem2* gene expression, VSG mice had a decreased serum level of sTREM2 (**Fig. 3C**), suggesting a reduced cleavage of membrane-bound TREM2 in macrophages due to improved inflammation^21^. Next, we performed differential gene expression analysis between LAMs from VSG, Sham AL, and Sham PF mice. Compared with LAMs from Sham AL (37 DEGs), and to a lesser extent Sham PF (24 DEGs), LAMs from VSG mice showed an increased expression of genes involved in lysosomal activity (*Lyz2*, *Ctsl, Ctss,* and *H2-Eb1*), antigen presentation (*H2-Eb1*), repression of inflammation (*Egr1*), and fatty acid metabolism (*Lipa*) (**Supplemental Table 4** and **Fig. 3D**). In contrast, several genes associated with inflammation (*Cd83*, *Nfkb1*, *Junb*, and *Mmp7*) were downregulated in LAMs from VSG mice (**Fig. 3D**). Pathway analysis revealed an upregulation of pathways associated with immune activation and lysosomal activity such as “Chemokine signaling”, “Cytokine receptor interaction”, and “Phagosome” in LAMs from VSG mice (**Fig. 3E**). Similarly, gene set enrichment analysis showed that genes involved in “Chemokine signaling”, “Lysosome”, “Peroxisome” and “Fatty acid metabolism” had increased expression in VSG LAMs, compared with Sham PF controls (**Fig. 3G**). Overall, these data highlight lysosomal and metabolic regulatory mechanisms by which VSG may sustain the protective function of LAMs against MASH in response to VSG.

**Figure 3.**
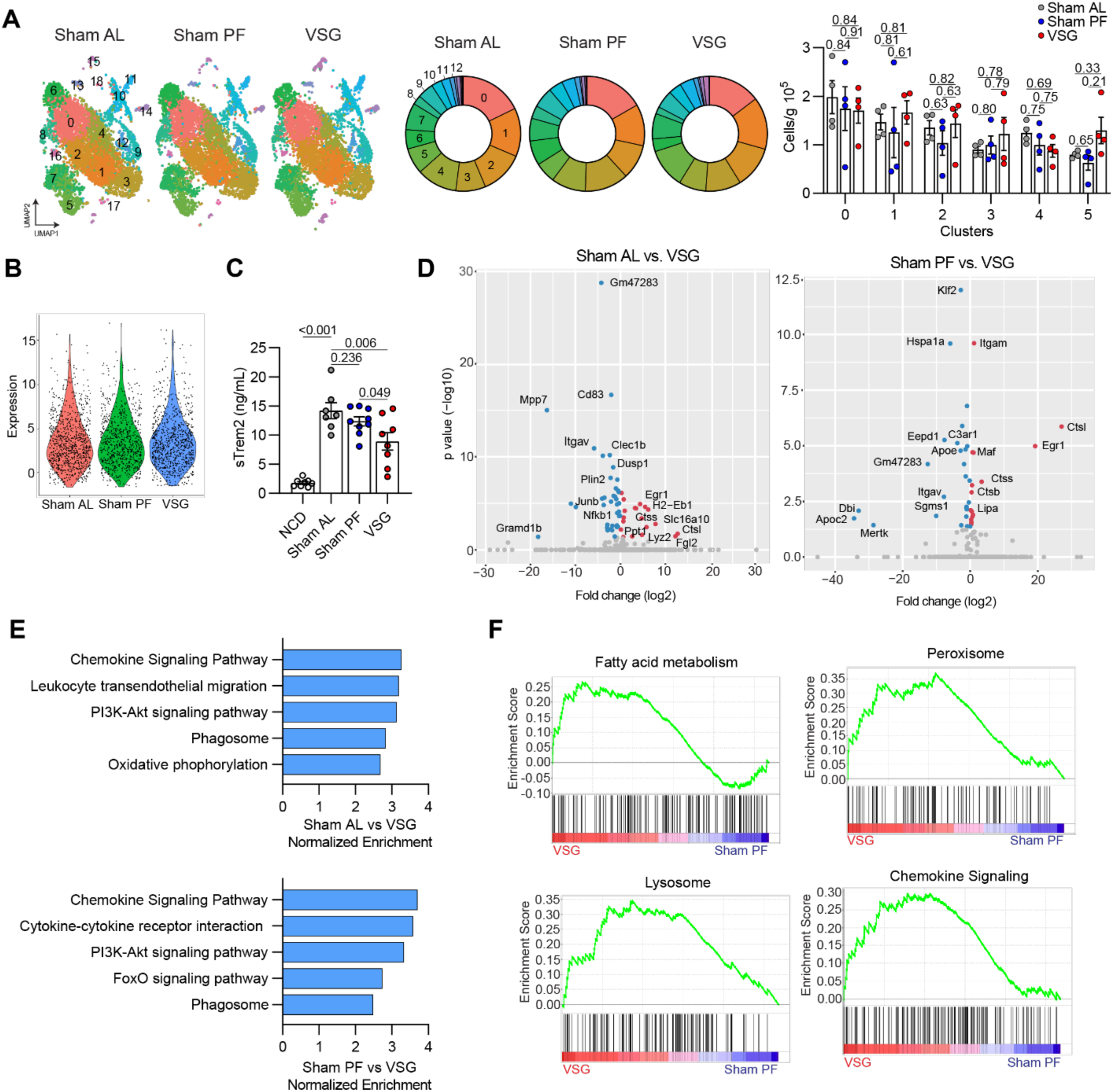
VSG enhances lipid metabolism and lysosomal gene programs in hepatic LAMs. (**A**) UMAPs (left), relative cluster proportion (middle), and number of cells per cluster (right) and (**B**) Expression of *Trem2* in cluster 1 (LAMs) from the scRNA-seq analysis of monocytes and macrophages from Sham AL, Sham PF, or VSG groups. (**C**) Concentration of soluble TREM2 (sTREM2) in the serum of NCD (n = 8), Sham AL (n = 7), Sham PF (n = 9), and VSG (n = 8) mice 5 weeks post-surgeries. (**D**) Volcano plots showing differentially expressed genes (DEGs) between Sham AL and VSG (left), and Sham PF and VSG (right) LAMs, (**E**) Pathway analysis of DEGs in LAMs from Sham AL vs. VSG (top) and Sham PF vs. VSG (bottom) comparisons, and (**F**) Gene seat enrichment analysis (GSEA) of the DEGs between Sham PF and VSG LAMs in the scRNA-seq data. The cell number and sTREM2 concentration data were analyzed by one-way ANOVA with Holm-Šídák multiple comparison test. Differential expression testing was performed by a Wilcoxon rank-sum test. Pathway analysis was performed by Generally Applicable Gene-set Enrichment Analysis (GAGE, p-value < 0.1). Data are biological experimental units presented as mean ± SEM.

### Hepatic TREM2^+^ LAMs mediate the reparative effects of VSG against MASH

Recent work has shown that systemic TREM2 deficiency worsens diet-induced MASH as TREM2 is required for LAM survival, the metabolic coordination between LAMs and hepatocytes, and the clearance of dying liver cells ^21, 22, 23, 40^. To determine if LAMs directly mediate the reparative process induced by VSG against MASH, we performed sham or VSG surgeries on WT and TREM2-deficient (TREM2 KO) mice fed an HFHC diet for 12 weeks. Expression of *Trem2* was not detectable in bone marrow-derived macrophages (BMDM) from TREM2 KO mice (**Extended Data Fig. 4A**). Five weeks post-surgery, WT and TREM2 KO mice showed a similar degree of weight loss (**Fig. 4A**) and an increase in fecal lipid excretion (**Fig. 4B**) in response to VSG. In agreement with our previous experiments, we found that sTREM2 was reduced following VSG in the serum of WT mice but was not detectable in TREM2 KO mice (**Extended Data Fig. 4B**).

**Figure 4.**
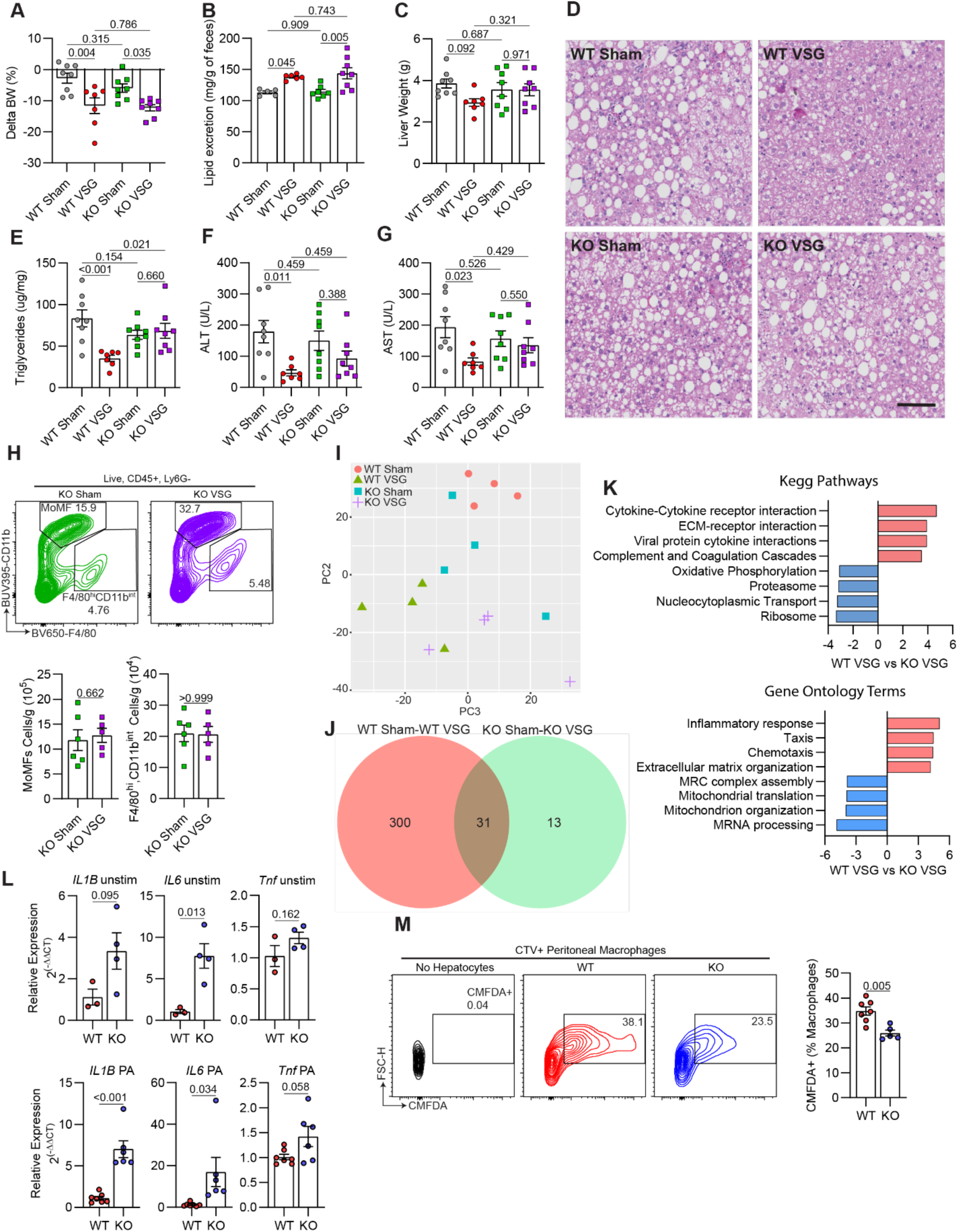
Hepatic TREM2^+^ LAMs mediate the reparative effects of VSG against MASH. (**A**) Body weight change post-surgery, (**B**) Fecal lipid content, (**C**) Liver weight, (**D**) Representative H&E-stained liver sections, (**E**) Hepatic triglyceride content, (**F**) Serum ALT and (**G**) AST, (**H**) Representative flow plot (top) and quantification of monocyte-derived (MoMFs) and F4/80^hi^ CD11b^int^ macrophages, (**I**) Unsupervised PCA of bulk RNA-seq gene expression data from F4/80^+^ sorted macrophages. (**J**) Venn diagram with the number of differentially expressed genes (DEGs) between Sham and VSG in WT and TREM2 KO macrophages, and (**K**) Pathway analysis (KEGG and Gene ontology) of DEGs between WT and TREM2 KO macrophages from sham or VSG mice. C57BL6/J (WT) and TREM2 KO (KO) mice were fed an HFHC diet for 12 weeks, assigned to either sham or VSG surgeries, and analyzed 5 weeks post-surgery (WT Sham, n = 5-8; WT VSG, n = 6-7; KO Sham, n = 5-8; and KO VSG, n = 5-8). (**L**) Gene expression of inflammatory cytokines in bone marrow-derived macrophages from WT (n = 3-7) or TREM2 KO (n = 4-6) mice left unstimulated (unstim, top) or stimulated with palmitate (PA, bottom). (**M**) Representative flow plots (left) and quantification (right) of CellTracker Green (CMFDA)-positive macrophages following coculture of peritoneal macrophages from WT (n = 7) and TREM2 KO (n = 5) mice with CMFDA-labeled apoptotic AML12 hepatocytes. Data from four experimental groups were analyzed by one-way ANOVA with Holm-Šídák multiple comparison test. Pathway analysis was performed by GAGE (p-value < 0.05). Data from two experimental groups were analyzed by Mann Whitney tests. Data are biological experimental units presented as mean ± SEM.

Notably, while VSG ameliorated MASH progression in WT mice, this effect was blunted in TREM2 KO mice (**Fig. 4C-G**). Compared with their Sham controls, VSG failed to decrease the liver weight (**Fig. 4C**), hepatic steatosis (**Fig. 4D and 4E**), ALT (**Fig. 4F**), and AST (**Fig. 4G**) in TREM2 KO mice, suggesting that TREM2 is required for the VSG-induced reversal of MASH. To explore the underlying mechanisms, we first assessed whether TREM2 deficiency alters the hepatic macrophage populations in mice with MASH before and after VSG. We found no differences in the number of MoMFs, emKCs, moKCs, and VSIG4- macrophages between WT and TREM2 KO fed the HFHC diet for 12 weeks without any intervention (**Extended Data Fig. 4C**). Similarly, the blunted effect of VSG in TREM2 KO mice was not associated with alterations in the number of hepatic macrophage subsets (**Fig. 4H and Extended Data Fig. 4D**). To determine the potential mechanisms by which TREM2 KO mice are resistant to the beneficial effects of VSG, we magnetically sorted and performed bulk RNA sequencing on total macrophages from the livers of WT and TREM2 KO mice after sham or VSG surgeries. Unsupervised PCA showed a distinct separation between WT Sham and WT VSG, whereas there was no distinction between TREM2 KO Sham and TREM2 KO VSG macrophages (**Fig. 4I**). Differential gene expression analysis revealed a more robust response of WT macrophages to VSG (331 DEGs), compared with that of TREM2 KO cells (44 DEGs) (**Fig. 4J and Extended Data Table 5**). In agreement, unsupervised clustering of the top 500 most variable genes revealed substantial gene expression differences and clustering between WT Sham and VSG macrophages but not between TREM2 KO Sham and VSG cells (**Extended Data Fig. 4E**). We performed pathway and gene ontology (GO) analyses and found that TREM2 KO macrophages from VSG mice upregulated genes enriched in inflammatory and immune activation pathways such as “cytokine-cytokine receptor interaction” and the GO terms “inflammatory response” and “chemotaxis” (**Fig. 4K**), suggesting that TREM2 is required for preventing an inflammatory activation of macrophages. To test this possibility, we examined the response of BMDMs from WT and TREM2 KO mice with or without stimulation with palmitate (PA) in vitro to mimic the lipid-rich environment of the MASH liver. Compared with WT controls, TREM2 KO BMDMs showed a markedly increased expression of *Il1b*, *Il6*, and *Tnf*, regardless of stimulation (**Fig. 4L**). Next, to determine whether TREM2 facilitates the ability of macrophages to clear apoptotic cells, we induced apoptosis in AML12 hepatocytes by PA treatment (**Extended Data Fig. 4F**) and co-cultured them with either WT or TREM2 KO peritoneal macrophages. Consistent with its key role in efferocytosis, TREM2 was required for macrophages to perform effective efferocytosis of apoptotic hepatocytes (**Fig. 4M**). Together, these data suggest that hepatic LAMs mediate the VSG-induced reversal of MASH by repressing inflammation and facilitating efferocytosis in a TREM2-dependent manner.

### VSG increases the content of inflammatory lipid species in hepatic macrophages

Our scRNA-seq data shows that hepatic LAMs are uniquely equipped with the enzymatic machinery to recognize, scavenge, and catabolize lipids^20, 38^. Because our data showed that total liver TGs decreased in response to VSG while hepatic LAMs upregulate lipid metabolism genes, we performed metabolic profiling of sorted F4/80^+^ macrophages from Sham AL and VSG livers to determine how bariatric surgery alters their lipid content. We performed metabolite profiling by liquid chromatography-tandem mass spectrometry (MxP® Quant 500 kit, Biocrates), which revealed 317 unique detectable metabolites in macrophages. The majority of these were lipid species, predominantly TGs and phosphatidylcholines, although we also detected amino acids and bile acids (**Fig. 5A**). Unsupervised principal component analysis (PCA) of detected metabolites showed a moderate clustering of VSG samples with more separation among Sham AL specimens (**Fig. 5B**). We found no difference in the concentration of total TGs in the mcarophages from Sham AL and VSG mice (**Fig. 5C**). However, when we performed chain length enrichment analysis of all lipid species, we found that macrophages from Sham AL mice were enriched in species with longer chain lengths while those from VSG mice were enriched in species of shorter chain length (**Fig. 5D**). Quantification of the major lipid families showed that macrophages from VSG mice had increased total phosphatidylcholines and sphingolipids, but no changes in total, cholesterol esters, fatty acids, glycosylceramides, ceramides, sphingolipids, and diacylglycerols (**Fig. 5E**). To further assess the effects of VSG on the lipid profile of hepatic macrophages, we assessed the composition of the individual lipid species. VSG macrophages had increased levels of phosphatidylcholines particularly species containing 2 acyl-bound (aa), one acyl- and one alkyl-bound (ae), and monounsaturated fatty acids (**Fig. 5F and Extended Data Fig. 5**). The role of phosphatidylcholines in macrophages is unclear with studies reporting both pro-inflammatory^41^ and anti-inflammatory^42^ responses. VSG macrophages were also enriched in sphingolipids containing long, very long-chain fatty acids, and hydroxyl-free species (**Fig. 5G**), as well as ceramides rich in very long-chain fatty acids (**Fig. 5H**). Given that sphingolipids acylated with fatty acids give rise to ceramides^43^, and that excess ceramides have detrimental effects on the liver including steatosis^44^, insulin resistance^45^, inflammation, and oxidative stress^46^, it is possible that VSG macrophages protect the liver from their detrimental effects. Although there were trending increases in VSG macrophages, we were unable to detect significant differences in the content of mono- and poly-unsaturated fatty acids (**Fig. 5I**) and cholesterol esters (**Fig. 5J**), compared with Sham controls. As the liver has decreased steatosis after VSG, the increase of macrophage intracellular lipids suggests an improved ability to clear lipids after the surgery.

**Figure 5.**
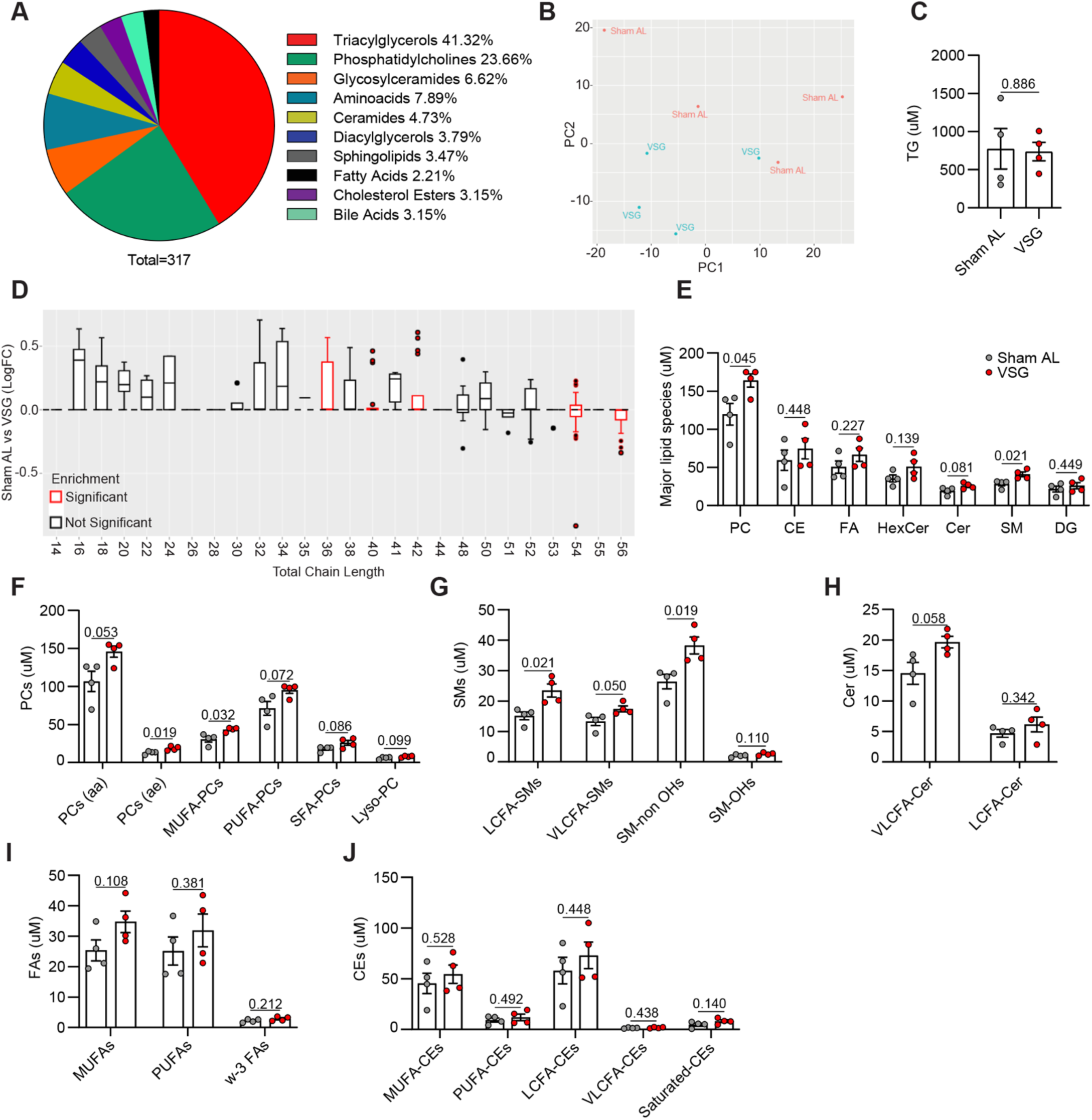
VSG increases the content of inflammatory lipid species in hepatic macrophages. (**A**) Composition of detectable metabolites in macrophages, (**B**) Unsupervised PCA of metabolite data, (**C**) Concentration of triacylglycerols (TG), (**D**) Chain length enrichment analysis of lipid species. The box represents the interquartile range, the middle line is the median, and the top and bottom lines indicate quartile 3 and quartile 1, respectively. Dots indicate outliers and red boxes indicate chain lengths with statistical significance between Sham AL and VSG, (**E**) Concentration of major lipid species including phosphatidylcholines (PC), cholesterol esters (CE), fatty acids (FA), glycosylceramides (HexCer), ceramides (Cer), sphingolipids (SM), and diacylglycerols (DG), (**F**) PC subspecies such as PCs (aa), PCs (ae), monounsaturated fatty acid (MUFA) PCs, polyunsaturated fatty acid (PUFA) PCs, and saturated fatty acid (SFA) PCs, (**G**) SM subspecies including long chain fatty acid (LCFA) SMs, very long chain fatty acid (VLCFA) SMs, and SMs with or without hydroxyl (-OH) groups, (**H**) Cer subspecies including VLCFA-Cer and LCFA-Cer, (**I**) FA subspecies including MUFAs, PUFAs, and omega-3 (w-3) FAs, and (**J**) CE subspecies determined by mass spectrometry and liquid chromatography of hepatic macrophages isolated from the livers of HFHC-fed mice assigned to Sham AL (n = 4) or VSG (n = 4) 5 weeks post-surgery. Data were analyzed by Welch’s two-sided t tests. Data are biological experimental units presented as mean ± SEM.

### Spatial transcriptomic of MASH livers following bariatric surgery reveal an improved metabolic status in the macrophage microenvironment

Given that our scRNA-seq analysis revealed substantial changes in the gene expression profile of hepatic macrophages, we explored the effects of VSG on the hepatic areas surrounding these cells using spatial transcriptomic analysis of liver sections (Nanostring GeoMx). Tissues from NCD, Sham AL, Sham PF, and VSG mice were collected 5 weeks after surgery and stained with a fluorescently labeled antibody against the macrophage marker CD68 to capture gene expression changes in CD68^-^ microanatomic areas (**Fig. 6A**). Following imaging and sequencing, the initial data was subjected to quality checks, filtering, and scaling, leading to a total of 7074 detectable genes and 32-36 regions of interest (ROI) per group (**Fig. 6B**) Unsupervised PCA of the normalized genes showed a substantial separation between CD68-expressing ROIs, and to a lesser extent between experimental groups (**Fig. 6C**). Due to the small size of their ROIs, we were unable to detect meaningful gene expression data from the CD68^+^ macrophage areas. However, differential expression gene analysis of the CD68^-^ ROIs revealed 185, 103, and 139 DEGs (FDR<0.05, FC > 1.5; **Supplemental File 6**) between NCD vs. Sham AL (**Fig. 6D**), Sham AL vs. and VSG (**Fig. 6E**), and Sham PF vs. VSG (**Fig. 6F**), respectively. To better understand the biological meaning of these changes, we performed enrichment and pathway analyses. Gene set enrichment analysis revealed that our analysis covered a wide range of cellular pathways including “metabolism”, “immune system”, “signal transduction” and “metabolism of proteins” (**Extended Data Fig. 6A**). Compared with the NCD group, pathway analysis showed that Sham AL mice had downregulated genes involved in the “metabolism of steroids” and “respiratory electron transport” and upregulation of “plasma lipoprotein remodeling” and “chemokine receptors bind chemokines” (**Fig. 6G**). Compared with Sham AL, VSG downregulated pathways involved in “biological oxidations”, “metabolism of amino acids”, and “metabolism of lipids” in agreement with the reduced lipid accumulation observed in VSG mice. On the other hand, VSG upregulated pathways such as “degranulation of neutrophils/platelets” and “complement cascade” indicative of an active immune and tissue remodeling response (**Fig. 6H**). Compared with the Sham PF group, we observed a downregulation of metabolic pathways and an upregulation of complement pathways in the VSG ROIs, similar to the VSG-induced changes relative to the Sham AL group (**Fig. 6I**). Overall, these findings highlight the VSG-induced transcriptomic changes in metabolic and immune pathways in the microenvironment surrounding hepatic macrophages that are associated with MASH reversal.

**Figure 6.**
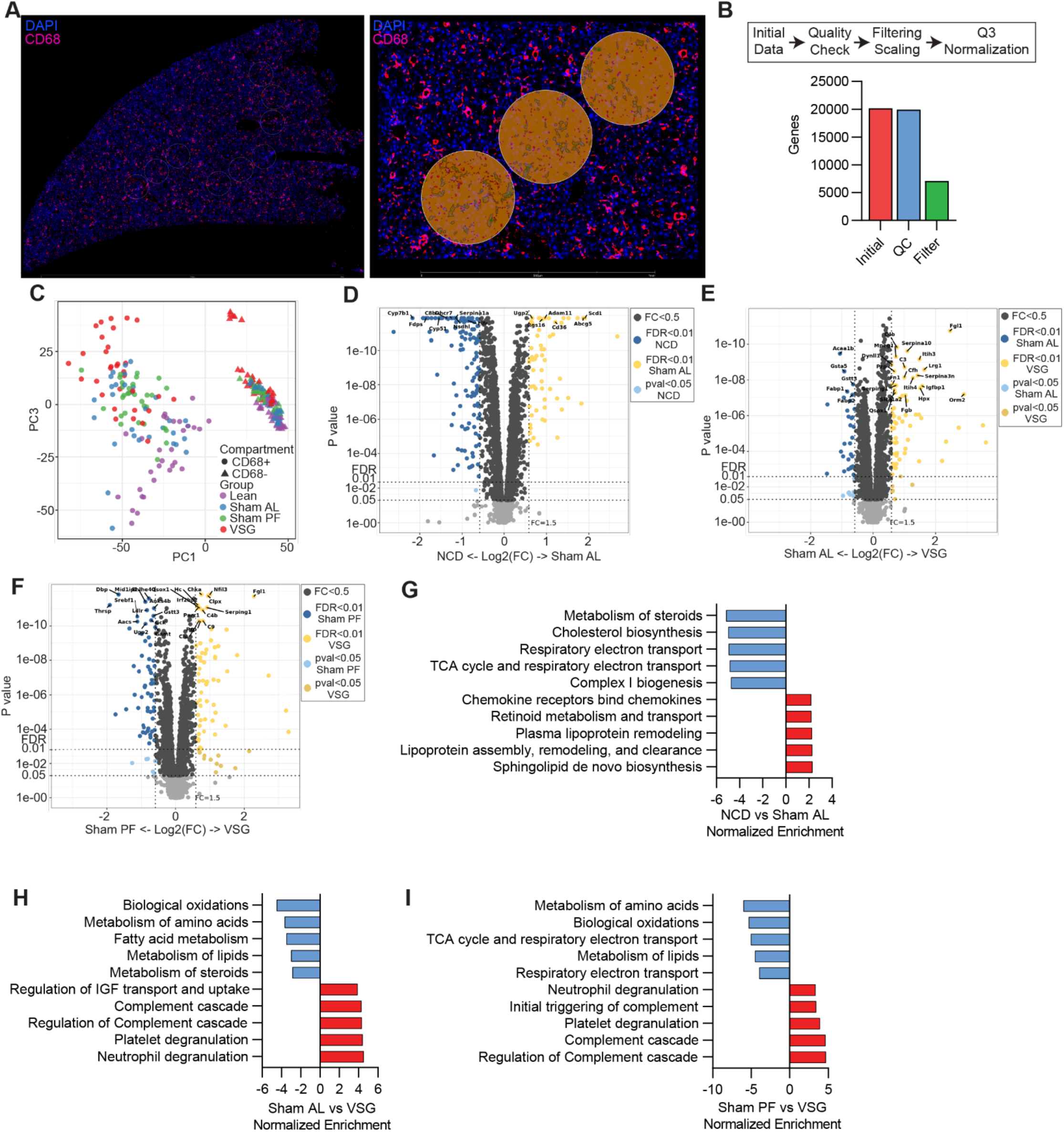
Spatial transcriptomic of MASH livers following bariatric surgery reveals an improved metabolic status in the macrophage microenvironment. (**A**) Representative immunofluorescence images showing an entire section (left) and a magnified field (right) with CD68^-^ (orange) and CD68^+^ regions of interest (ROIs, green) in liver specimens analyzed by spatial transcriptomics (Geomx). (**B**) Steps in data processing (top) and number of genes after processing (bottom), (**C**) Unsupervised PCA of gene expression data from CD68^+^ and CD68^-^ ROIs, Volcano plots showing differentially expressed genes between (**D**) NCD and Sham AL, (**E**) Sham AL and VSG, (**F**) Sham PF and VSG CD68^-^ ROIs. Pathway analysis between (**G**) NCD and Sham AL, (**H**) Sham AL and VSG (**I**) Sham PF and VSG CD68^-^ ROIs. Liver specimens were collected from C57BL6/J (WT) fed either a NCD or an HFHC diet for 12 weeks, assigned to Sham AL, Sham PF, or VSG surgeries, and analyzed 5 weeks post-surgery (n = 4). Data were analyzed utilizing a Mann-Whitney test and corrected with a Benjamini-Hochberg procedure. Pathway analysis was performed by GSEA (p-value < 0.05).

Finally, we determined whether our main findings could be relevant to human patients with MASH undergoing VSG. We performed spatial transcriptomics on needle biopsy specimens from a patient with MASH, collected before and one year after VSG (**Extended Data Fig. 6B**) as VSG resulted in the resolution of MASH (NAS 4 to NAS 0 after 12 months, **Table 1**). We also included a liver specimen from a donor patient with MASH without any surgical intervention. After staining with cytokeratin (green) and the macrophage marker CD68 (magenta), 32 ROIs were annotated to define hepatic zones I, II, and III (**Extended Data Fig. 2C**). Unsupervised PCA of detectable genes revealed a substantial separation between specimens but small differences between hepatic zones (**Extended Data Fig. 2D**). Pathway analysis of these comparisons revealed several downregulated pathways involved in the “metabolism of lipids”, “cholesterol biosynthesis”, and “metabolism of steroids” (**Extended Data Fig. 2E**), consistent with the MASH resolution observed in this patient. To explore if human LAMs are responsive to VSG as in our mouse studies, we assessed the expression of genes involved in the LAM differentiation program including *Trem2*, *Plin2*, *Ctss*, *Cd36*, and *Lipa*^38, 40^. LAM genes were primarily enriched in zone I of the liver (**Extended Data Fig. 2F)**, in agreement with a recently published spatial transcriptomic study^20^. Furthermore, the LAM genes were upregulated in the Pre-VSG and MASH samples but were lower in the Post-VSG sample (**Extended Data Fig. 2F**). These data suggest that LAMs correlate with disease progression in human MASH.

**Table 1.** Human spatial transcriptomics specimen information.

| Specimen | Bx Date | NAS Score | Age | Sex | Race | BMI | DM Dx | MS Dx | Alcohol Use |
| --- | --- | --- | --- | --- | --- | --- | --- | --- | --- |
| PreVSG | 11/13/19 | 4 | 42 | F | Caucasian | 35.3 | PreDM | No | None |
| PostVSG | 12/23/20 | 0 | 43 | F | Caucasian | 26.7 | None | No | None |
| NASH | N/A | 6 | 70 | F | Caucasian | N/A | N/A | N/A | N/A |

## DISCUSSION

Bariatric surgeries are the most effective options available for the treatment of obesity complications including MASLD and MASH^7, 8, 9, 47^. Indeed, a recent randomized trial showed that bariatric surgery is more effective than lifestyle intervention and medical care in the treatment of patients with MASH, including those with liver fibrosis^48^. The beneficial effects of bariatric surgeries against obesity-related diseases are usually attributed to the mechanical restriction of the stomach and malabsorption of nutrients. However, accumulating evidence shows that the surgeries exert profound effects in the regulation of several physiological functions ranging from caloric intake^49^, respiration^50^, intestinal bile acids and lipid absorption^30^, iron metabolism^17^, and immune responses^51^. Here we show that VSG induces weight-loss-independent improvements in MASH in a process that requires TREM2-expressing macrophages. These data implicate the innate immune system in the control of inflammation as an important mechanism of VSG action. Previous work has shown that systemic inflammatory mediators associated with obesity gradually decline following bariatric surgery in humans^52^ and mouse models of disease^53, 54^. As TREM2- deficient mice do not respond to the surgery, our findings further demonstrate that the reduced inflammation in the liver is not secondary to caloric restriction or weight loss but a contributing factor.

Recent research has shown that MASH is associated with the emergence of LAMs as a specialized subset of macrophages that are required for adequate metabolic coordination, lipid handling, extracellular matrix remodeling, and clearance of apoptotic hepatocytes^21, 22, 23, 40, 55^. Notably, metabolic stress and inflammation cause TREM2 shedding in LAMs resulting in the loss of immune homeostasis and aggravated disease^21^. Here we show that TREM2 is necessary for the VSG-induced reversal of disease in a process associated with decreased shedding of TREM2. Mechanistically, we show that TREM2 is required for dampening hepatic inflammation and effective efferocytosis of lipid-laden hepatocytes by macrophages. Furthermore, these two processes are likely dependent on each other as efferocytosis has been shown to suppress inflammation via the liver X receptor^56^. Given that TREM2 is a master regulator of the LAM phenotype^36^, we reasoned that TREM2 would mediate their reparative role induced by VSG. Indeed, our findings indicate that VSG restores several TREM2-dependent pathways such as “AKT-PI3K signaling”, required for effective suppression of inflammation^57^, and “oxidative phosphorylation”, which is typical of macrophages in anti-inflammatory or reparative states^58^. Overall, our data show that VSG induces a restorative function in macrophages in a TREM2- dependent manner.

Given that the local micro-environment has a profound impact on the transcriptional program and function of macrophages and neighboring cells^58^, we used spatial transcriptomics to assess the effects of VSG on hepatic macrophage-adjacent areas. In agreement with reduced steatosis, lipid metabolism pathways were suppressed by VSG in the non-macrophage areas. However, one of the main immune pathways that was upregulated following VSG was the complement system. This was surprising as increased levels of complement proteins have been associated with MASH severity in humans^59^ and complement has been shown to increase hepatic de novo lipogenesis, inflammation, and insulin resistance^60^. Additional studies are needed to determine if complement proteins play a role in the response of the MASH liver to VSG due to the limited biological implications that can be drawn from transcriptomic data. Nevertheless, hepatic complement proteins have been shown to promote a reparative or apoptotic cell-clearing phenotype^61^. In support of this notion, we found that macrophages from VSG mice have an enrichment of several lipid species at the same time as the liver presents with decreased steatosis and a downregulation of genes associated with lipid metabolism.

Not surprisingly, bariatric surgery induces rapid and profound changes in the gut microbiota likely due to anatomical and metabolic mechanisms^62, 63^. Increased *Lactobacillus*, considered a beneficial commensal bacteria^64, 65^, has been proposed to be a mechanism by which VSG improves obesity and glucose tolerance^66^. As the potential ligands of TREM2 include bacterial products^67^, it is possible that gut-derived microbial products may have direct effects on hepatic LAMs or other macrophage subsets through pattern recognition receptors^68^, or gut-derived metabolites drained via the portal vein^69^. For example, increased intestinal butyrate may be an additional mechanism by which VSG induces a reparative macrophage phenotype as VSG restores the levels of butyrate-producing *Lactobacillus*^70^.

One limitation of our study is that we utilized a mouse with a systemic TREM2 deficiency. Although TREM2 is primarily expressed by myeloid cells, bacterial and viral infections have been reported to induce TREM2 expression in CD4 T cells^71, 72^. Additionally, TREM2-expressing macrophages in the adipose tissue have been shown to prevent obesity and adiposity, which could influence the development of hepatic steatosis^36^. While several key experiments such as our scRNAseq and spatial transcriptomics analyses show that hepatic LAMs are responsive to VSG, we cannot rule out the potential contribution of adipose tissue LAMs to the VSG-induced protective effects in the liver.

In summary, here we show that VSG induces a reversal of MASH independent of weight loss accompanied by a substantial remodeling of the gut microbiota and hepatic macrophages. Here, we report that hepatic TREM2^+^ LAMs respond to VSG by increasing their expression of lysosomal and oxidative phosphorylation genes while downregulating inflammatory genes. Notably, TREM2 deficiency ablates the protective effects of VSG, providing causal evidence that LAMs mediate the beneficial effects of bariatric surgery against MASH.

## MATERIALS AND METHODS

### Animals

Five-week-old C57BL/6J (000664) and TREM2-deficient (TREM2 KO, 027197) male mice were purchased from The Jackson Laboratory and maintained in a pathogen-free, temperature-controlled environment. At six weeks of age, mice were fed a high-fat high-carbohydrate (HFHC, 40% kcal palm oil, 20% kcal fructose, and 2% cholesterol supplemented with 23.1 g/L d-fructose and 18.9 g/L d-glucose in the drinking water) diet ad-libitum for 12 weeks to induce MASH. Subsequently, they received either a sham operation or VSG surgery. Post-surgery, the mice were provided a liquid diet for 2 days and the HFHC diet was slowly re-introduced. 7 days post-surgery the mice were only fed the HFHC diet for the remainder of the study. To account for weight-loss-independent effects, there was a sham ad-libitum-fed group (Sham AL) and a sham group that was pair-fed (Sham PF) to the VSG group to match the weight loss post-surgery. The amount of HFHC diet fed to the Sham PF group was the mean consumption of the VSG group from the preceding day. Additionally, we included a group of mice that were fed a normal chow diet (NCD) for the duration of the study. All animal experiments were approved by the University of Minnesota Institutional Animal Care and Use Committee.

### Macrophage characterization and isolation

Intrahepatic immune cells were isolated from perfused livers using enzymatic digestion as previously described^73^. For flow cytometry, single cell suspensions were incubated with Zombie NIR (1:200, Biolegend) for 20 min at room temperature, TruStain FcX (1:50, Biolgend) for 5 min at room temperature, and with fluorophore-conjugated primary antibodies (1:100) for 30 min at 4°C. Following the staining, cells were washed and fixed with 100 μL Fixation Buffer (BioLegend) for 20 min at 4°C. Flow cytometry data were acquired on a Fortessa flow cytometer (BD Biosciences) and analyzed using Flowjo (TreeStar) software. For macrophage-specific assays, macrophages were isolated from immune cell suspensions using anti-F4/80 microbeads and a MACS separator (Miltenyi Biotech).

### Bone marrow-derived macrophages

Bone marrow-derived macrophages (BMDM) were generated as previously described^74^. Briefly, bone marrow was isolated from the femurs and tibias of male mice, then seeded in 2mL of conditioned media with macrophage colony-stimulating factor (mCSF, 25 ng/mL, BioLegend). Five days later, 1 mL of conditioned media with mCSF (50 ng/mL) was added. Seven days after seeding, the BMDMs were washed and stimulated with palmitate for 24 hours (200 uM, Cayman Chemical). Total RNA was extracted from the BMDMs using the RNeasy Plus Mini kit (Qiagen). cDNA was prepared using the iScript cDNA Synthesis kit (Bio-Rad). Gene expression was calculated using the 2(-ΔΔCT) method and normalized to GAPDH.

### Efferocytosis assessment

Peritoneal macrophages were isolated from WT and TREM2 KO mice by lavage with phosphate-buffered saline (PBS) with 2mM EDTA, as previously described^75^. Macrophages were then plated on 24 well non-tissue culture-treated plates (Corning) in conditioned RPMI with 10% fetal bovine serum and 1% penicillin and streptomycin for 1 hour. The media was replaced to remove non-adherent cells and the cells were cultured for 4 hours. Subsequently, the peritoneal macrophages were stained with CellTrace Violet (CTV, ThermoFisher) for 20 min at 37°C. AML12 hepatocytes (ATCC) were treated with PA (1200 μM) for 24 hours to induce apoptosis (**Extended Data Fig. 4F)**, then stained with CellTracker Green (CMFDA) for 30 min at 37°C. Following, CTV+ macrophages were cocultured with CMFDA+ apoptotic hepatocytes at a ratio of 1:4, respectively, for 2 hours. The cells were then lifted, centrifuged, and resuspended for flow cytometry acquisition. Flow cytometry data were acquired on a Fortessa flow cytometer (BD Biosciences) and analyzed using Flowjo (TreeStar) software. Efferocytosis capacity was calculated as a percentage of CTV+, and CMFDA+ cells relative to total CTV+ cells.

### Fecal lipid extraction

Fecal lipid extraction was performed as previously described^76^. Feces were collected, added to 5 mL of PBS, and vortexed until the fecal pellets disintegrated. Next, 5 mL of chloroform:methanol (2:1, v/v) was added, vortexed, and centrifuged at 1,000g for 10 min. The lower liquid phase was collected, transferred to a glass tube, and evaporated in a fume hood. The remaining lipids were weighed.

### MASH phenotyping

Hepatic triglycerides were assessed using a calorimetric assay (Cayman Chemical). Concentrations of serum ALT and AST were assessed by the University of Minnesota Veterinary Medical Center’s Clinical Pathology Laboratory. Liver sections were fixed in 10% formalin and hematoxylin & eosin (H&E) and Picro Sirius Red staining was performed by the Biorepository & Laboratory Services at the UMN Clinical and Translational Science Institute. The collagen area was assessed using Picro Sirius Red stained liver sections as previously described^77^. NAS scoring of the H&E-stained liver sections was performed by a blinded pathologist.

### Metabolite assessments

Metabolite assays were performed by the University of Minnesota’s Center for Metabolomics and Proteomics. Hepatic macrophage metabolites were determined utilizing Biocrates MxP® Quant 500 kit. Lipids and hexoses were measured by flow injection analysis-tandem mass spectrometry (FIA-MS/MS) and small molecules were measured by liquid chromatography-tandem mass spectrometry (LC-MS/MS) using multiple reaction monitoring (MRM) for the detection of analytes utilizing a Shimadzu LC-20AD XR (Shimadzu USA Manufacturing Inc.) coupled to a Sciex QTRAP 5500 mass spectrometer (Sciex,). Hepatic portal vein bile acids and short chain fatty acids were acquired using an Agilent 1290 series HPLC (Agilent,) with an Acquity C18 BEH (2.1mm x 50mm, 1.7 um) column coupled to an Agilent 6495C Triple Quadrupole (Agilent) mass spectrometer in negative ion-mode. Bile acid samples were prepared by combining 100 uL of hepatic portal vein serum, 20 uL of the internal standard, 30 ul of HCL (1 M), and 1 mL of acetonitrile^78^. The acquired data was imported into Agilent MassHunter software for additional analysis. Hepatic portal vein short chain fatty acids samples were prepared by combining 90uL of hepatic portal vein serum 10 uL of the internal standard, and 420 μL of cold methanol. Concentrations of metabolites were calculated in the Biocrates MetIDQ™ software (Biocrates Life Sciences AG). Metabolites missing more than 20% of measurements were excluded from the statistical analysis. Missing values were imputed using the k-nearest neighbors approach as implemented in the *impute* R-package^79^. Hypothesis testing was performed using a Welch’s two sided t-test. Lipid set enrichment was performed using LipidSuite software^80^.

### Bulk RNA sequencing

Total RNA was extracted from hepatic macrophages using the RNeasy Plus Mini kit (Qiagen). Samples were sequenced on a Novaseq 6000 using a 150 PE flow cell at the University of Minnesota Genomics Center. The SMARTer Stranded RNA Pico Mammalian V2 kit (Takara Bio) was used to create Illumina sequencing libraries. Differential gene expression analysis was performed using edgeR (Bioconductor). Gene ontology and pathway analysis were completed using iDEP.96 software^81^.

### Fecal microbiota assessment

DNA was extracted from single fecal pellets using the PowerSoil DNA isolation kit (QIAGEN) on the automated QIAcube platform using the inhibitory removal technology (IRT) protocol. The V4 hypervariable region of the 16S rRNA gene was amplified and sequenced using the 505F/816R primer set^82^. Paired-end sequencing was done at a length of 301 nucleotides (nt) on the Illumina MiSeq platform by the University of Minnesota Genomics Center using their previously described dual-index method^83^. Raw sequence data are deposited in the Sequence Read Archive under BioProject accession number SRP325945. Sequence data processing and analysis were performed using Mothur ver. 1.41.1^84^. Sequences were aligned against the SILVA database^85^ (ver. 138) for processing. Clustering of operational taxonomic units was done at 99% similarity using the furthest-neighbor algorithm and taxonomic classifications were made against the Ribosomal Database Project (ver. 18)^86^. Compositional data are reported from non-normalized data. For statistical comparisons, samples were rarefied to 10,000 reads/sample^87^. Ordination of samples was done using principal coordinate analysis^88^, and differences in community composition were evaluated by analysis of similarity (ANOSIM), with Bonferroni correction for multiple comparisons^89^. Differences in alpha diversity were determined by ANOVA with Tukey’s posthoc test and differences in genera relative abundances were evaluated by the non-parametric Kruskal-Wallis test, with pairwise comparisons done using the Steel-Dwass-Critchlow-Fligner procedure. To determine dissimilarity from baseline composition, SourceTracker2^90^ was used with baseline samples in each group designated as the source. All statistics were evaluated at *α*=0.05 prior to correction for multiple comparisons.

### Spatial transcriptomics

Fixed moused liver sections were mounted and stained with DAPI and an anti-CD68 to create macrophage-free ROIs. Fixed human liver sections were mounted and stained with DAPI/anti- CD68/anti-panCK antibodies to create zone-specific ROIs. Spatial sequencing was performed as previously described using the GeoMx platform (NanoString)^91^. Analysis of the sequencing data was performed on the GeoMx Digital Spatial Profiler (NanoString).

### Human specimens information

We used human de-identified specimens previously collected by the Biological Materials Procurement Network (BioNet) at the University of Minnesota. We used specimens collected by liver biopsy from one patient before and one year VSG and from a patient diagnosed with MASH without any surgical intervention (**Table 1**). Samples were obtained with institutional review board approval.

### Single-cell RNA sequencing

Two million CD45^+^ and F4/80^+^ cells from each sample were individually incubated with cell multiplexing oligos (CMO) to allow for downstream multiplexing of samples. Cells were washed with PBS containing 0.04% BSA and centrifuged at 400g for 5 minutes at room temperature. The supernatant was carefully discarded, and the cell pellet was resuspended in 100 μL of CMO for 5 minutes. Cells were washed with 1.9mL PBS containing 1% BSA and centrifuged at 400g for 5 minutes at 4⁰C. Then, cells were washed three times using 2 mL of wash buffer. Following, samples were combined at a concentration of 1,500 cells/μL and loaded into capture ports of a 10x-Genomics chip following a single cell 3’ kit. Separate libraries were created for gene expression and CMO then sequenced utilizing a Novaseq S4 chip (2 x 150bp PE). Feature quantification was performed using cellranger (version 3.0, 10X Genomics). All subsequent analyses were performed using Seurat 3.1.1^92^. After sample demultiplexing of CMOs, we retained single cells and normalized within each pool using the 4 sctransform method in Seurat^92^. Visualization of different clusters was enabled using Uniform Manifold Approximation and Projection (UMAP) dimensional reduction^93^. The Seurat cell ID function was utilized to identify monocytes and macrophages, which were re-clustered for analysis. Differential expression testing was performed using the default Wilcoxon rank-sum test. To evaluate the developmental path of macrophages, we inferred trajectories using slingshot^37^. To limit our analyses to the developmental stages of interest, we first created a data set only containing the major monocyte and macrophage clusters. Then, we inferred the trajectories with cluster 6 set as the origin and cluster 1 as the terminal state. To visualize gene expression patterns across developmental pseudotime and across lineages, we first fit a negative binomial generalized additive model to the observed gene expression, which was then used to predict smoothed expression values for each cell and lineage using the tradeseq package^94^. All Seurat, slingshot, and tradeseq analyses were conducted in R (v.4.1.0). Pathway analysis was completed using iDEP.96^81^ and DAVID^95^ software. Gene set enrichment was performed using the GSEA software^96^.

### Statistical analysis

Statistical significance was determined using a one-way ANOVA with Holm-Šídák multiple comparison test using GraphPad Prism (version 10.0.2). All data analyzed by ANOVA followed a Gaussian distribution (Shapiro-Wilk test) and had equal variances (Brown-Forsythe). For datasets that failed the normality test, statistical significance was determined using multiple non-parametric two-sided Mann-Whitney tests. For in vitro experiments, each data point represents a biological replicate with cells from one mouse. Mouse studies were repeated in at least two independent experiments. Data are presented as means ± SEM with statistical significance denoted.

## ACKNOWLEDGMENTS

This study was supported by the National Institute of Diabetes and Digestive and Kidney Diseases (DK122056 to X.S.R.) and the National Heart, Lung, and Blood Institute (R01HL155993 to X.S.R.). G.F. is supported by a T32 NIH training grant (T32DK083250) We recognize the staff from the Research Animal Resources, University Flow Cytometry Resource, Center for Metabolomics and Proteomics, Genomics Center, and the Clinical and Translational Science Institute at the University of Minnesota for their assistance.

## AUTHOR CONTRIBUTIONS

X.S.R., G.F., and S.I conceived the study and designed experiments. G.F. and X.S.R interpreted results, generated figures and tables, and wrote the manuscript. G.F., K.F., F.B, K.D., H.W., P.P., C.J., C.S., and R.A. performed experiments. O.A. analyzed and provided data from liver specimens. A.H. performed the analysis of single-cell RNA sequencing data. A.B., C.J., J.W., and D.G.M. provided feedback and supervised aspects of the study. X.S.R obtained funding for, supervised, and led the overall execution of the study.

---

**Extended Data Figure 1 (related to Figure 1).**
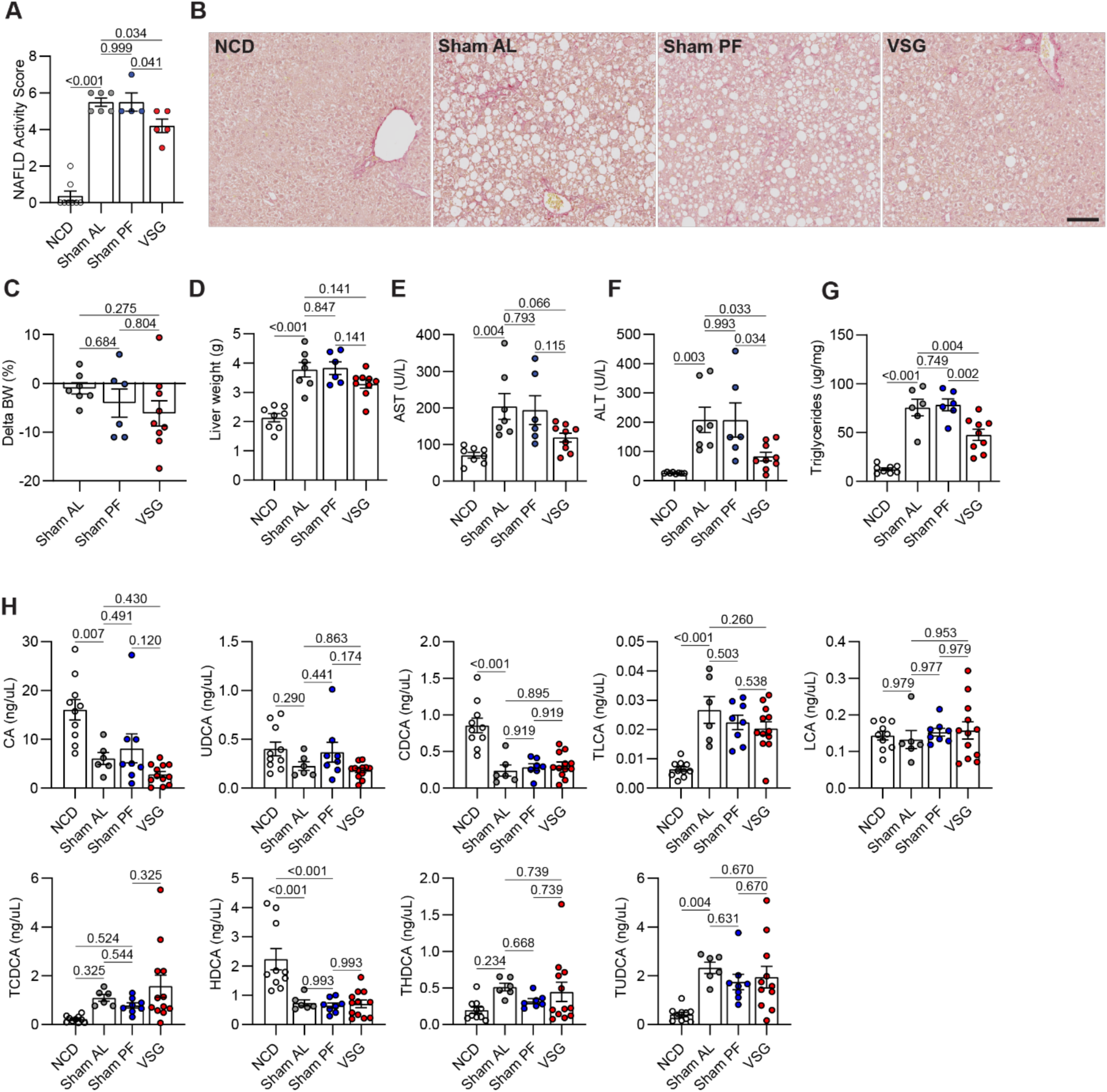
(**A**) NAFLD activity score; NCD (n = 8), Sham AL (n = 6), Sham PF (n = 4), VSG (n = 5), and (**B**) Representative Picro Sirius red-stained liver sections 5 weeks post surgeries. (**C**) Percentage change in body weight, (**D**) Liver weight, (**E**) Serum AST, (**F**) Serum ALT, and (**G**) Hepatic triglyceride content in C57BL6/J (WT) mice fed either an NCD (n = 8) for 22 weeks or an HFHC diet for 12 weeks, assigned to Sham AL (n = 6-7), Sham PF (n = 6), or VSG (n = 9) surgeries, and maintained on the HFHC diet for 10 weeks post-surgery. (**H**) Concentration of portal vein bile acids 5 weeks post-surgery: cholic acid (CA), ursodeoxycholic acid (UDCA), chenodeoxycholic acid (CDCA), taurolithocholic acid (TLCA), lithocholic acid (LCA), taurochenodeoxycholic acid (TCDCA), hyodeoxycholic acid (HDCA), taurohyodeoxycholic acid (THDCA), tauroursodeoxycholic acid (TUDCA); NCD (n = 10), Sham AL (n = 6), Sham PF (n = 8), VSG (n = 12). Data were analyzed by one-way ANOVA with Holm-Šídák multiple comparison test. Data are biological experimental units presented as mean ± SEM.

**Extended Data Figure 2 (related to Figure 2).**
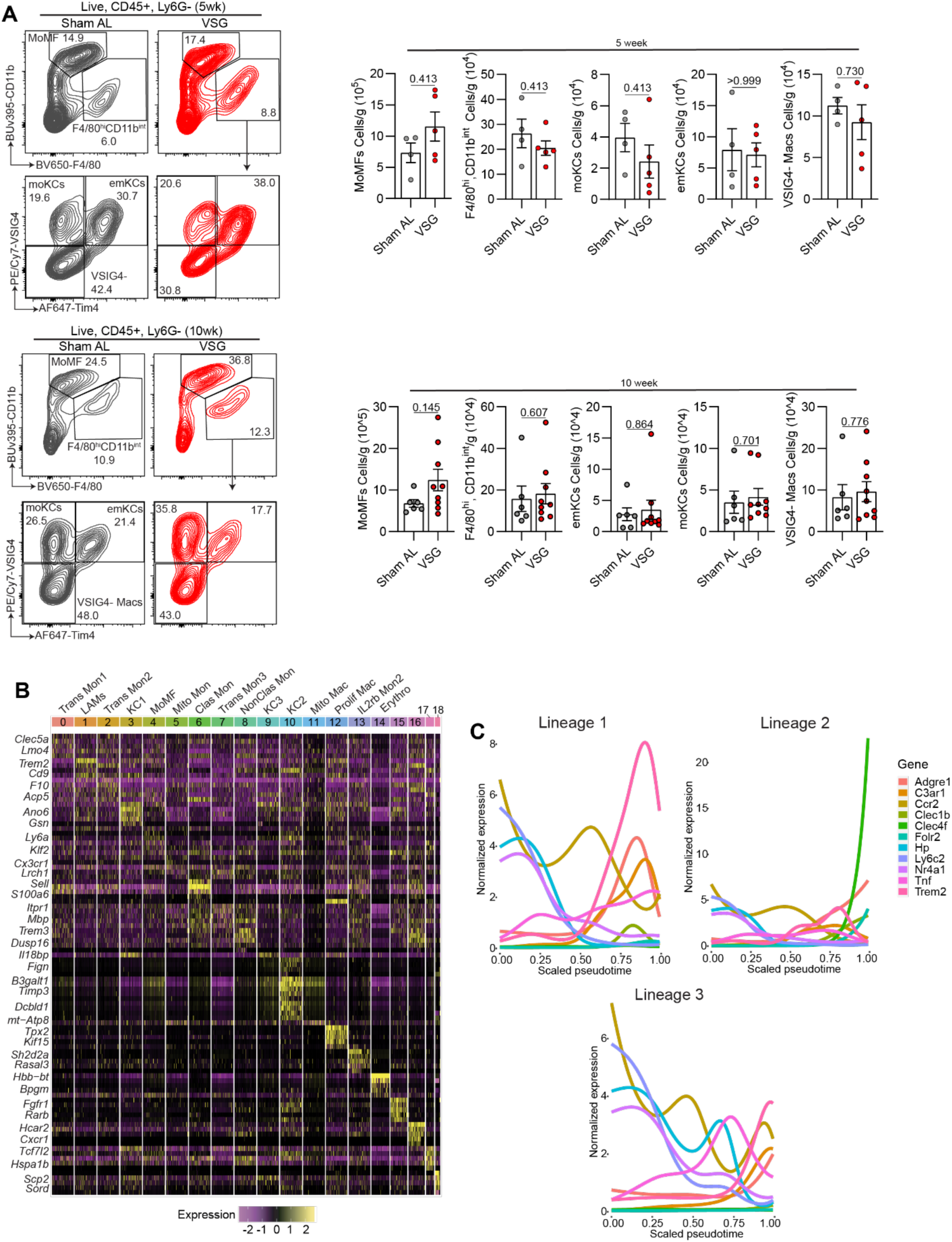
(**A**) Representative flow plots (left) and quantification of hepatic macrophage subsets per gram of liver (right) at 5 weeks (top) and 10 weeks post-surgery (bottom); 5 week Sham AL (n = 4), 5 week VSG (n = 5), 10 week Sham AL (n = 6) and 10 week VSG (n = 9). (**B**) Heat map of top genes for each cluster. (**C**) Gene expression over pseudotime for trajectories 1 (top left), 2 (top right) and 3 (bottom). Data were analyzed by Mann Whitney tests. Data are biological experimental units presented as mean ± SEM.

**Extended Data Figure 3 (related to Figure 3).**
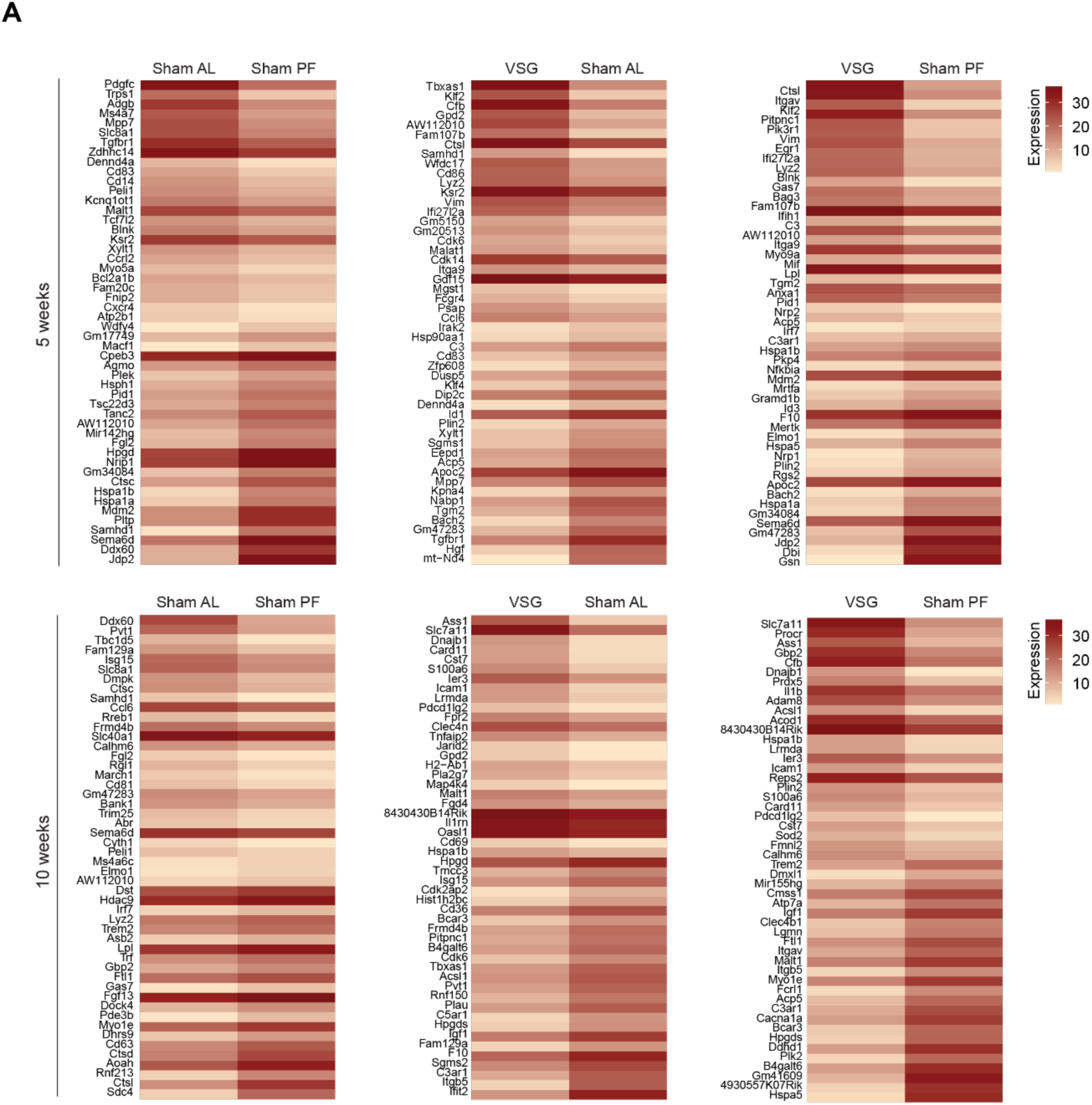
(**A**) Heat maps of top DEGs between all groups at both time points for total macrophages. All data presented in this figure (n = 4).

**Extended Data Figure 4 (related to Figure 4).**
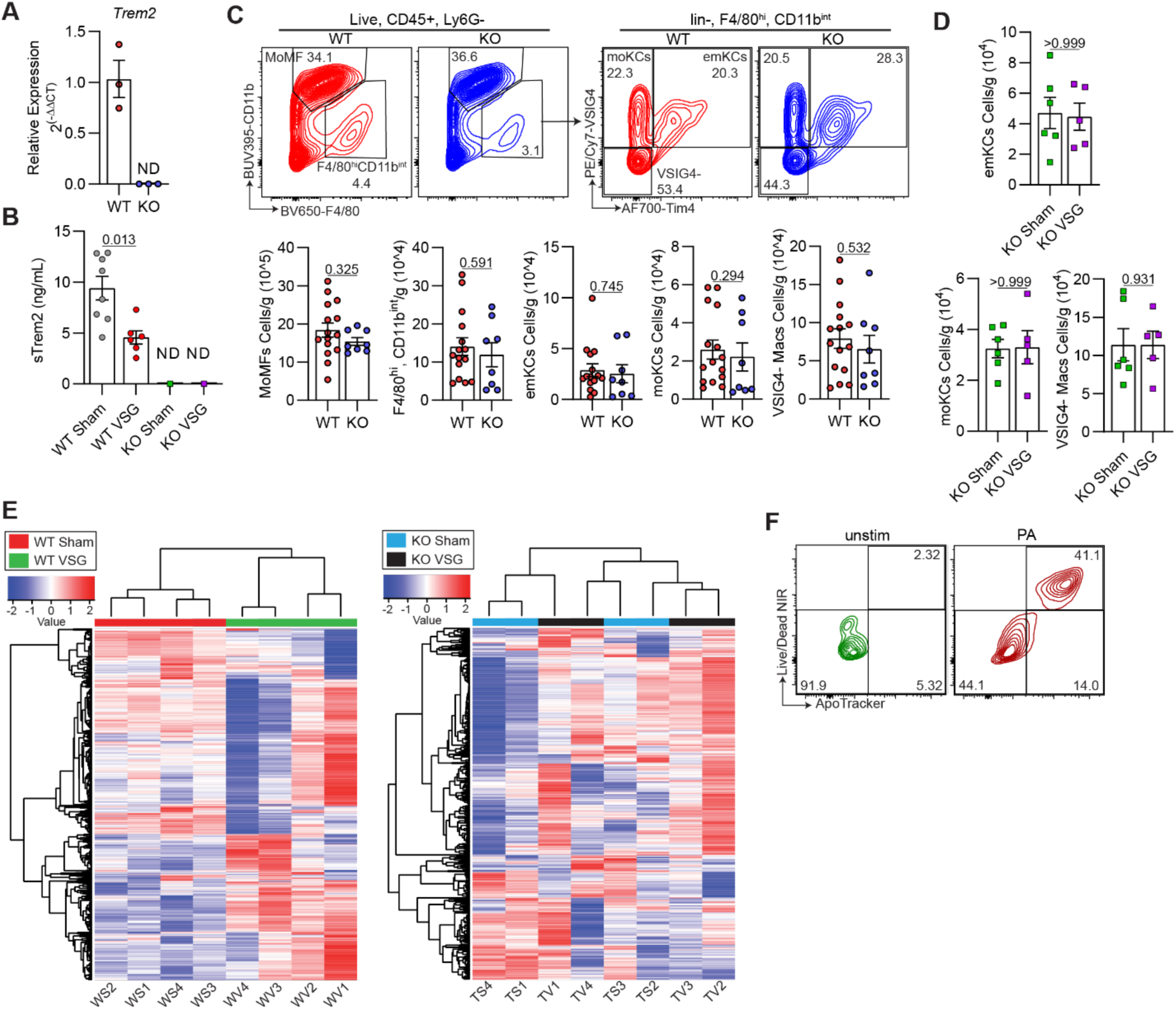
(**A**) Relative expression *Trem2* in WT and TREM2 KO BMDMs; WT (n = 3) and KO (n = 3). (**B**) Concentration of sTREM2 in the serum 5 weeks post-VSG; WT Sham (n = 8), WT VSG (n = 6), KO Sham (n = 8), and KO VSG (n = 8). (**C**) Representative flow plots (top) and quantification of hepatic macrophage subsets per gram of liver (bottom) after 12 weeks of HFHC feeding; WT (n = 15) and KO (n = 8). (**D**) Quantification of hepatic macrophage subsets per gram of liver 5 weeks post-VSG; WT (n = 5) and KO (n = 5). (**E**) Heat map of the top 500 most variable genes between WT Sham and WT VSG macrophages (left) and KO Sham and KO VSG macrophages (right). (**F**) Representative flow plots of AML12 cells that were either unstimulated or stimulated with PA. Data were analyzed by Mann Whitney tests. Data are biological experimental units presented as mean ± SEM.

**Extended Data Figure 5 (related to Figure 5).**
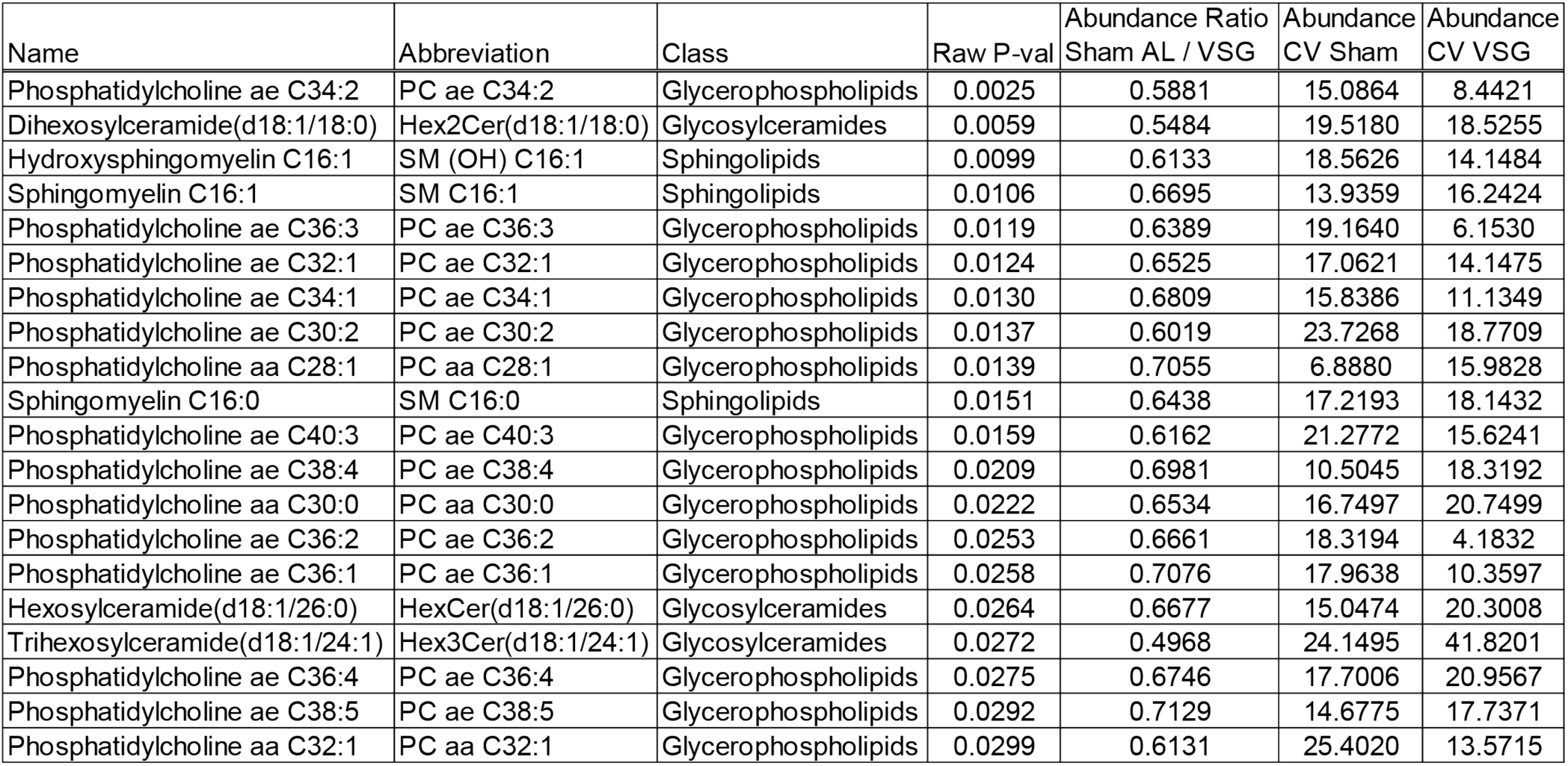
Table of top metabolites in hepatic macrophages 5 weeks post-VSG.

**Extended Data Figure 6 (related to Figure 6).**
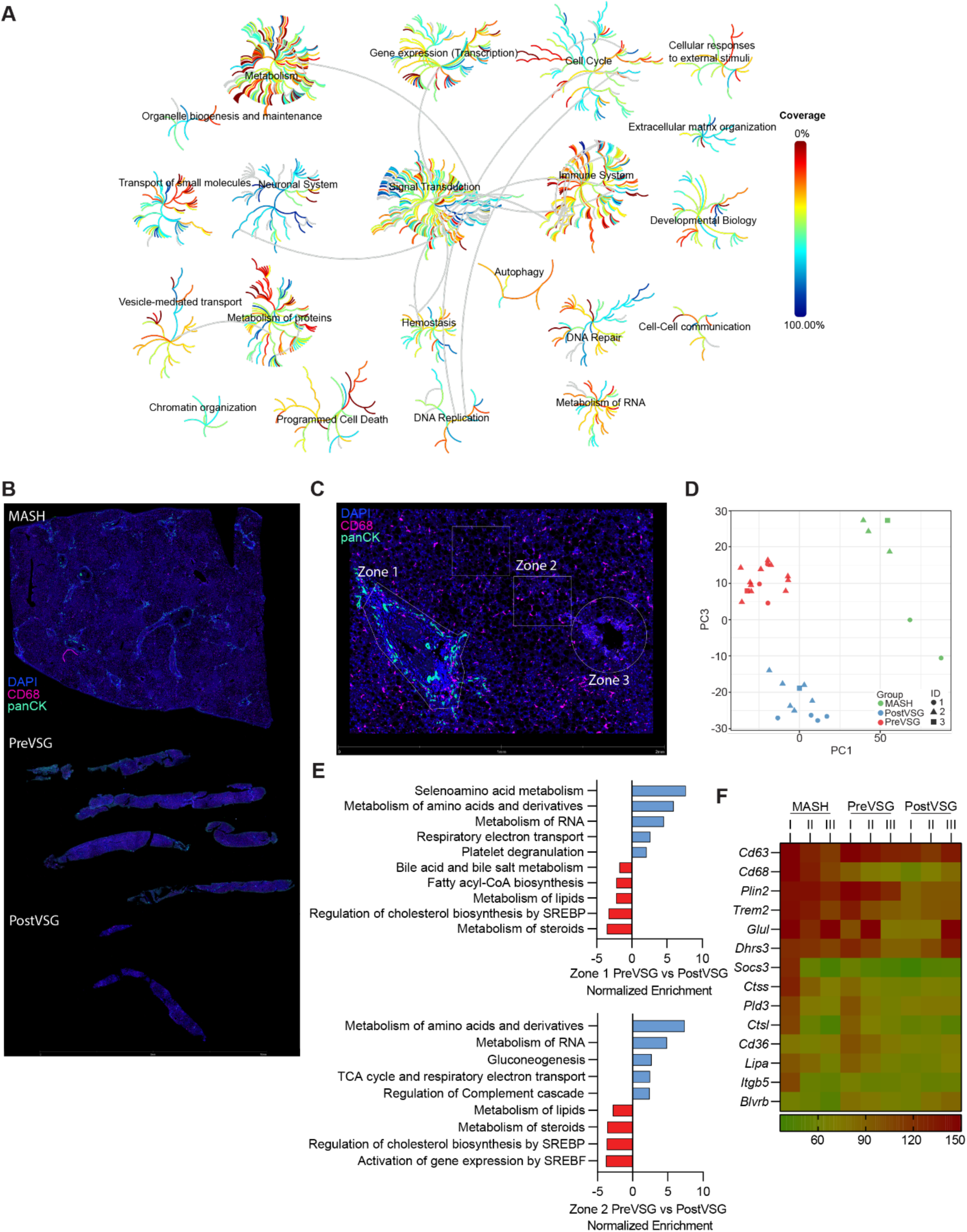
(**B-F**) Spatial sequencing of human liver samples pre-VSG and post-VSG. (**A**) Pathway coverage for mouse VSG GeoMx analysis. (**B**) Representative human liver sections used for GeoMx. (**C**) Representative human liver section used for GeoMx with liver zones indicated. (**D**) Unsupervised PCA of all human liver ROIs (n = 1). (**E**) Pathway analysis for Zone 1 PreVSG vs. Zone 1 PostVSG (top) and Zone 2 PreVSG vs. Zone 2 PostVSG (bottom). (**F**) Heatmap of LAM genes.

